# Comparative evaluation of glomerular morphometric techniques reveals differential technical artefacts between FSGS and normal glomeruli

**DOI:** 10.1101/2022.02.15.480420

**Authors:** John M Basgen, Anand C Reghuvaran, Qisheng Lin, Khadija Banu, Hongmei Shi, John Pell, Jenna DiRito, Gregory T Tietjen, Sudhir Perincheri, Dennis G Moledina, Francis Perry Wilson, Madhav C Menon

## Abstract

Morphometric estimates of mean glomerular volume (MGV) have clinical implications, over and above histologic data. However, MGV estimation is time-consuming, could waste tissue sections and requires expertise limiting its utility in retrospective clinical studies.

**Methods:** We evaluated MGV using both plastic and paraffin-embedded tissue from control and FSGS mice (n=10 each) using the gold-standard Disector/Cavalieri technique (Vglom-Cav) and other reported techniques [2- or 3-profile technique, Weibel-Gomez method (W-G)]. Within Vglom-Cav we examined the precision of MGV estimation while using MGVs obtained from 5- or 10-individual glomeruli measurements vs the true mean (20 glomeruli).

**Results:** In both FSGS and controls, we identified an acceptable precision of 10-glomerular sampling vs true MGV within Vglom-Cav technique [88 (79-94) % of MGV obtained were within 10% of the true MGV]. The 5-glomerular sampling was less precise [70 (56, 81) % of MGV obtained were within 10% of true MGV]. In plastic based techniques, 2- or 3-profile MGVs showed greater concordance with Vglom-Cav, than W-G MGV. The new 3-profile technique offered incremental benefit to the existing 2-profile method (improved Lin’s concordance in control and FSGS animals). We observed a consistent reduction of Vglom values within control animals (52+/-0.06%) in paraffin-embedded tissue (vs corresponding methods in plastic) demonstrating a clear shrinkage artefact due to tissue processing. FSGS glomeruli showed significantly less and more variable shrinkage artefact likely due to glomerular fibrosis.

**Conclusion:** We report the precision of 5- or 10-glomerular sampling for MGV estimation using controls and FSGS animals. We demonstrate and quantify the shrinkage bias in MGV during tissue processing for paraffin-embedding that also differentiated control animals and FSGS. Our findings have implications for experimental studies using glomerular morphometry.

## Introduction

Quantitative glomerular structural evaluation by morphometry has consistently revealed additional prognostic benefit over histology in human or experimental, diabetic-^1,2^, or non-diabetic-glomerular disease ^2–9^. For instance, we and others reported that FSGS was associated with larger glomerular volumes (Vglom) and podocyte hypertrophy, and that larger Vgloms predicted outcomes in human nephrotic syndromes^4,10^. Pathophysiologically, Vglom may reflect antecedent nephron injury and loss, with the consequent need for remaining nephrons (and glomeruli) to hypertrophy to increase the effective filtration surface area. Excessive or disproportionate glomerular hypertrophy (with respect to podocyte hypertrophy) promoted podocyte loss, glomerulosclerosis and progressive CKD^3,7^. Thus, Vglom estimation has diagnostic and prognostic implications.

The technical gold standard technique for Vglom estimation is the Disector/Cavalieri method^11^ using the Cavalieri equation (Vglom-Cav)^12,13^. This method is independent of glomerular shape, and able to measure individual glomerular volumes (IGV) as well as the mean Vglom (MGV) while providing a distribution of Vgloms within and across each individual. Vglom-Cav can be performed in plastic- or paraffin-embedded tissue (the latter requiring special ultramicrotomes), requires careful sectioning, can include only complete glomeruli and is time intensive. While being a highly precise technique for morphometry in dedicated nephrectomy studies^8,9^, its use in retrospective needle biopsy studies is limited. Other methods of MGV measurement applied to paraffin embedded tissue, which assume a spherical glomerular shape and could suffer formalin fixation artefacts, such as the two-profile method^13^, or Weibel Gomez^14^ (W-G) are more applicable for retrospective studies, albeit less precise.

A typical paraffin-embedded renal biopsy archived for retrospective studies includes limited numbers of glomeruli, from which further smaller sections are divided for plastic embedding. In recent data^15^ we reported that a subset of paraffin-embedded biopsy tissue from the NEPTUNE consortium^16^ had a mean 14+/− 10 (median 12) glomerular profiles. Prior data suggests that using Vglom-Cav from >10 glomeruli was not associated with significant improvements in the Coefficient of variation while calculating MGV^11^. Whether these sample sizes are also applicable to paraffin-based methods is not clear from current literature. Similarly, whether less time intensive morphometric techniques performed in plastic embedded tissue offer comparable precision to the Vglom-Cav is also unknown. A quantification of whether and how structural glomerular pathology (e.g., FSGS) influences MGV measurements by paraffin- or plastic-methods has also not been performed before.

Based on the significance of Vglom measurements to glomerular disease, we undertook a detailed evaluation of Vglom techniques, comparing each to Vglom-Cav using normal control mice and mice from FSGS models. We aimed to evaluate the number of glomeruli vs their precision in reflecting MGV by applying a modeling strategy that assumes random sampling of glomeruli to the gold-standard Vglom-Cav measurements. We quantify formalin fixation and paraffin embedding induced “shrinkage artefact” due to sample preparation that is present regardless of the individual techniques tested. Furthermore, the presence of FSGS reduced this shrinkage artefact albeit non-uniformly, possibly reflecting the variable presence of glomerular fibrosis in FSGS kidneys. The data here are informative for future experimental morphometric studies

## Results

### Samples used for Study

We restricted our analyses to Balbc/J mice to avoid any strain-related variation in comparisons, and used young male adult BALBc/J as controls (2-6 months age) for evaluating baseline morphometry (n=10). To evaluate morphometry during significant glomerular injury, we used mice from two FSGS models performed in BALBc/J (n=10) - hypertrophic injury and a previously published ageing model [See methods^10,17^]. The mice in these FSGS models had confirmed podocyte loss, glomerulosclerosis, proteinuria and azotemia^10,17^. In each animal, we evaluated three plastic-based morphometric techniques and three paraffin-based methods, comparing each with Vglom-Cav. Within the gold standard Vglom-Cav method, we evaluated sample size using up to 20 individual glomeruli per animal, to generate the gold standard Vglom-Cav. We modified the previously described 2-profile technique^13^, and evaluated the inclusion of a third glomerular profile using an additional sequential tissue section to MGV calculation (ie 3-profile technique; see methods). A summary of the paraffin- and plastic-morphometric techniques we used in this study; their important relative advantages are illustrated in Figure-1. For instance, like Vglom-Cav, the 2- or 3-profile techniques provided IGV and MGV, while W-G method could only provide MGV. Further, the latter three techniques assume spherical shape of glomeruli (incorporating a bias), while Vglom-Cav is shapeindependent.

**Figure-1:**
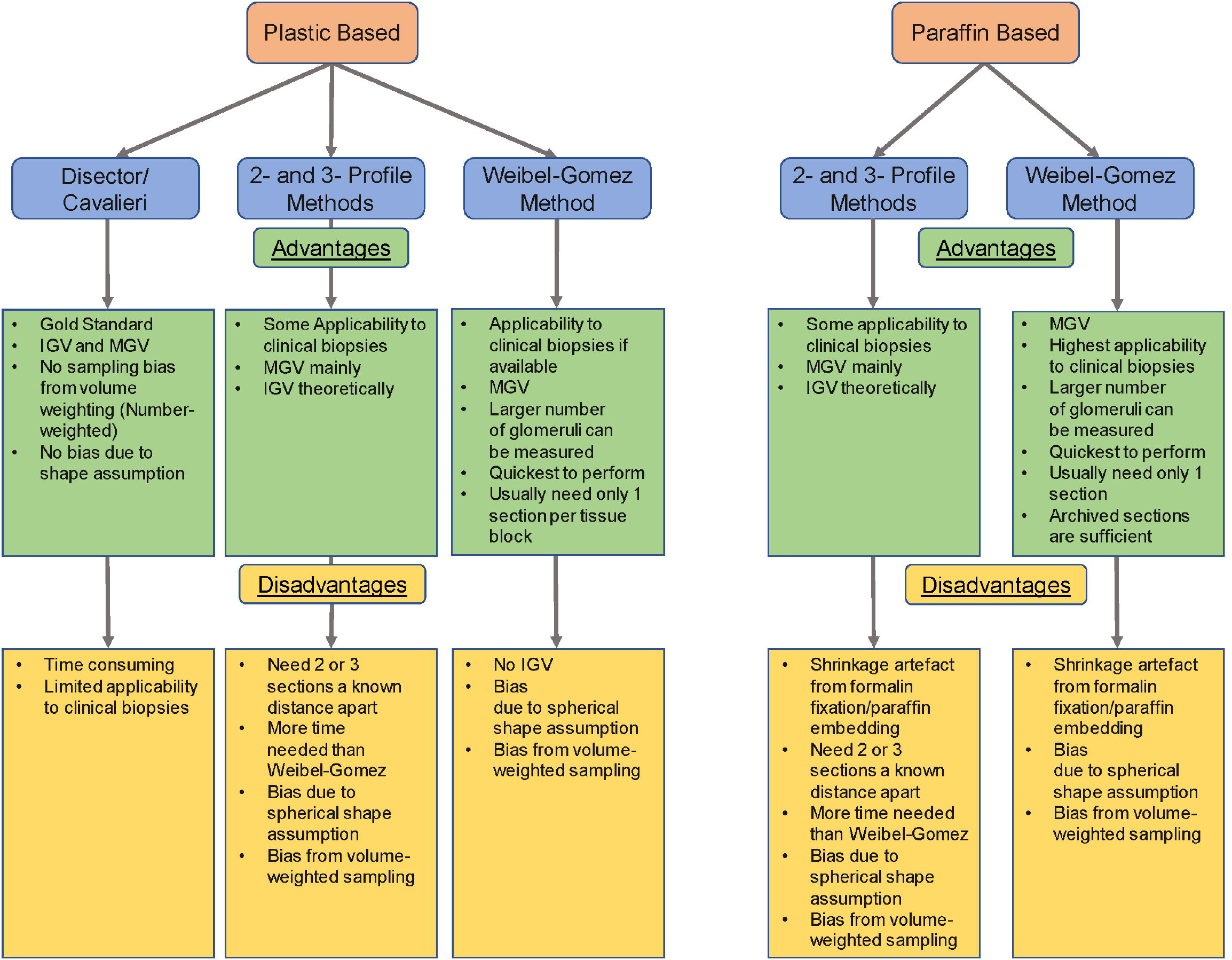
Plastic-embedded and Paraffin-embedded Morphometric techniques used in the study. Flow diagram tabulates the four plastic- and three paraffin-based techniques evaluated in our overall dataset of 10-control and 10-FSGS animals. The principal relative advantages and disadvantages of each technique within each tissue-embedding method are listed. MGV calculated from 20 individual glomeruli using the Disector/Cavalieri method is considered the gold-standard in these analyses to which other techniques were compared.

#### Empiric validation of calculating mean volume from a 5- or 10-glomeruli sample

Using Vglom-Cav either on plastic or paraffin tissue, prior work reported lack of significant additional benefit in MGV estimation above 10 glomeruli (individual Vgloms) sampled per specimen ^13,18^. Here, we empirically tested the hypothesis that mean volume obtained by sampling 5 or 10 glomeruli closely matches the true mean (i.e., that obtained by sampling 20 glomeruli). We used individual glomerular volumes (IGV) and calculated MGV from 1000 iterations of random sampling of 5 or 10 glomeruli amongst 20 glomeruli. In modelling 5-glomerular samples, amongst FSGS mice we noted that sample mean was within 5% of true mean only 42% of the time, similar to control animals (Table-1). When sampling 10 glomeruli 88 (79-94) % of the mean volumes obtained were within 10% of the true mean, 99.993 (99.94-99.992) % were within 25% of true mean, whereas only 64 (49-76) % were within 5% of the true mean. There were no significant differences in these proportions obtained when evaluating FSGS (n=5) or control (n=9) animals separately. Hence using our mouse models, these data provide a quantitative estimate of precision for MGV obtained from 10 IGVs when using Vglom-Cav.

**Table 1:** Proportion of times mean volume obtained by sampling 5 or 10 glomeruli was within given range of that obtained by sampling 20 glomeruli.

| Criteria | Sampling 5 glomeruli | Sampling 10 glomeruli |
| --- | --- | --- |
| **Overall (n=14)** | **Proportion (95% CI) within given % of true mean** |  |
| 5% | 42 (32, 53) % | 64 (49-76) % |
| 10% | 70 (56, 81) % | 88 (79-94) % |
| 25% | 97 (94, 98) % | 99.993 (99.94-99.9992) % |
| **FSGS (n=5)** |  |  |
| 5% | 42 (32, 53) % | 61 (40-79) % |
| 10% | 70 (56, 81) % | 89 (71-97) % |
| 25% | 97 (94, 98) % | 99.98 (99.7-99.999) % |
| **Control (n=9)** |  |  |
| 5% | 42 (32, 53) % | 65 (42-83) % |
| 10% | 70 (56, 81) % | 88 (72-95) % |
| 25% | 97 (94, 98) % | 100% |

*1000 iteration of random sampling of 5 or 10 glomeruli. Only includes those with 20 glomeruli. True mean=mean glomerular volume obtained by sampling 20 glomeruli.*

### Comparison of MGV measurements in plastic-embedded tissue in Control and FSGS mice

Since the above data suggested that in Vglom-Cav technique, 10 glomeruli provided a reasonable estimate of MGV as obtained from 20 glomeruli, we evaluated the relationships of MGV obtained by other plastic-based methods (Fig-1) with Vglom-Cav. Table-2a shows Lin’s concordance values between MGVs obtained by each technique compared to Vglom-Cav. In these analyses, concordance between MGV by Weibel Gomez was markedly lower than obtained by all other plastic based techniques. By each technique evaluated, differences in image magnifications (40X vs 100X) did not consistently or beneficially alter obtained concordance values. Furthermore, applying our novel 3-profile technique (modification of the 2-profile technique; See methods) showed an incremental but small improvement in Lin’s concordance vs the 2-profile method in control animals.

**Table 2A:**
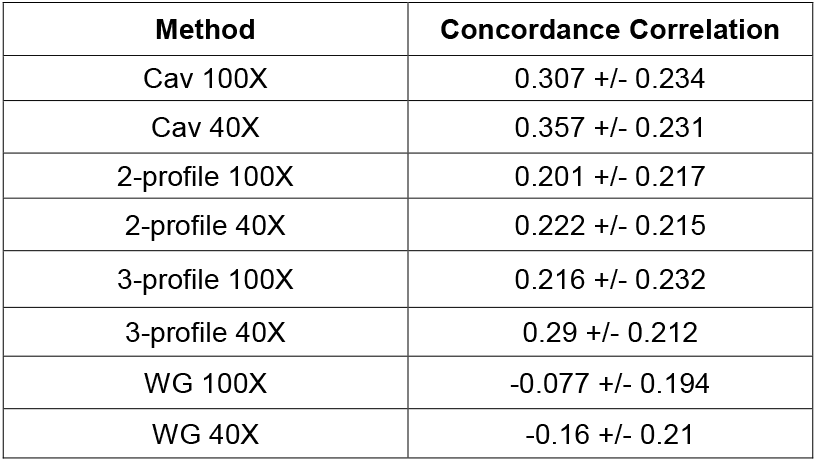
Lin’s concordance (SE) compared to gold standard mean volume in Control mice (N=10). All methods reflect the mean of 10 sampled glomeruli in plastic.

| Method | Concordance Correlation |
| --- | --- |
| Cav 100X | 0.307 +/- 0.234 |
| Cav 40X | 0.357 +/- 0.231 |
| 2-profile 100X | 0.201 +/- 0.217 |
| 2-profile 40X | 0.222 +/- 0.215 |
| 3-profile 100X | 0.216 +/- 0.232 |
| 3-profile 40X | 0.29 +/- 0.212 |
| WG 100X | -0.077 +/- 0.194 |
| WG 40X | -0.16 +/- 0.21 |

We then evaluated concordance between MGVs obtained by other plastic based techniques in FSGS animals, vs gold standard Vglom-Cav (shown in Table 2B). Here too we restricted our evaluation to MGVs obtained from 10 IGVs or glomerular profiles (for W-G method). Except in W-G method, a high concordance was observed in MGVs in Cav, 2- or 3-profile method when compared to the gold standard. Similar to observations in control animals, increased image magnification for measurements had inconsistent benefit to MGV by all techniques, while 3-profile technique again showed a limited and incremental benefit in concordance values over 2-profile technique.

**Table 2B:** Lin’s concordance (SE) compared to gold standard mean volume in FSGS mice (N=6). Al methods reflect the mean of 10 sampled glomeruli in plastic.

| Method | Concordance Correlation |
| --- | --- |
| Cav 100X | 0.988 +/- 0.011 |
| Cav 40X | 0.987 +/- 0.013 |
| 2-profile 100X | 0.691 +/- 0.250 |
| 2-profile 40X | 0.751 +/- 0.209 |
| 3-profile 100X | 0.789 +/- 0.170 |
| 3-profile 40X | 0.800 +/- 0.166 |
| WG 100X | 0.085 +/- 0.489 |
| WG 40X | 0.108 +/- 0.481 |

We then developed regression equations to relate MGVs obtained from 10 glomeruli using Cavalieri, 2- or 3-profile or W-G techniques to the Vglom-Cav gold standard (Table-2C).

**Table 2C:** Linear regression equations relating various methods (10 glom, plastic) to the gold standard.

| Method | Slope | Intercept |
| --- | --- | --- |
| Cav 100X | 0.989 | -2800 |
| Cav 40X | 0.946 | 6100 |
| 2-profile 100X | 0.707 | 52500 |
| 2-profile 40X | 0.784 | 42700 |
| 3-profile 100X | 0.661 | 54800 |
| 3-profile 40X | 0.668 | 54800 |
| WG 100X | 0.152 | 141100 |
| WG 40X | 0.17 | 138500 |

### Relating MGV from Paraffin-based techniques to Plastic-based techniques in Control and FSGS animals

While morphometry can be performed in plastic or paraffin embedded tissue, formalin-fixation and paraffin embedding are known to cause shrinkage with bias or distortion of morphometric measurements in other tissues^19,20^. Here we compared MGV measurements between 2- or 3-profile, and W-G performed in plastic vs these same techniques in paraffin tissue within each animal. In control animals, we identified a nearly uniform reduction in MGVs in paraffin based measurements vs corresponding plastic measurements (across all techniques mean+/− SD ratios = 0.52+/− 0.06; Mean of SDs across techniques =0.11) This was reflected in the uniform bias identified in Bland-Altman plots of controls evaluating paraffin:plastic MGV ratio(Figure 2A). These findings are consistent with a shrinkage artefact with tissue processing for paraffin embedding. Contrastingly, FSGS animals showed lower mean reductions in MGV compared to corresponding plastic measurements via each technique (mean+/− SD ratios = 0.78+/− 0.07; Mean of SDs across techniques =0.28). The greater variability in bias in FSGS animals is represented as a Bland Altman plot [Figure 2B]. Mean Shrinkage was also significantly different between control and FSGS animals [Fig 2C]. The variability of shrinkage *within each FSGS animal* when ratios were obtained from different techniques likely represented greater interglomerular IGV variation in kidneys with FSGS pathology. This was reflected in the coefficient of Variation in IGVs obtained in FSGS glomeruli as we previously reported^10^. Moreover, there was marked variability in the paraffin:plastic MGV ratios between animals with FSGS. As expected, Masson-trichrome stain of FSGS animals showed glomerular fibrosis, albeit of varying extent (Figure 2D). These data suggest glomerular fibrosis which develops in FSGS glomeruli could restrict the shrinkage artefact in paraffin-based MGV estimation.

**Figure-2:**
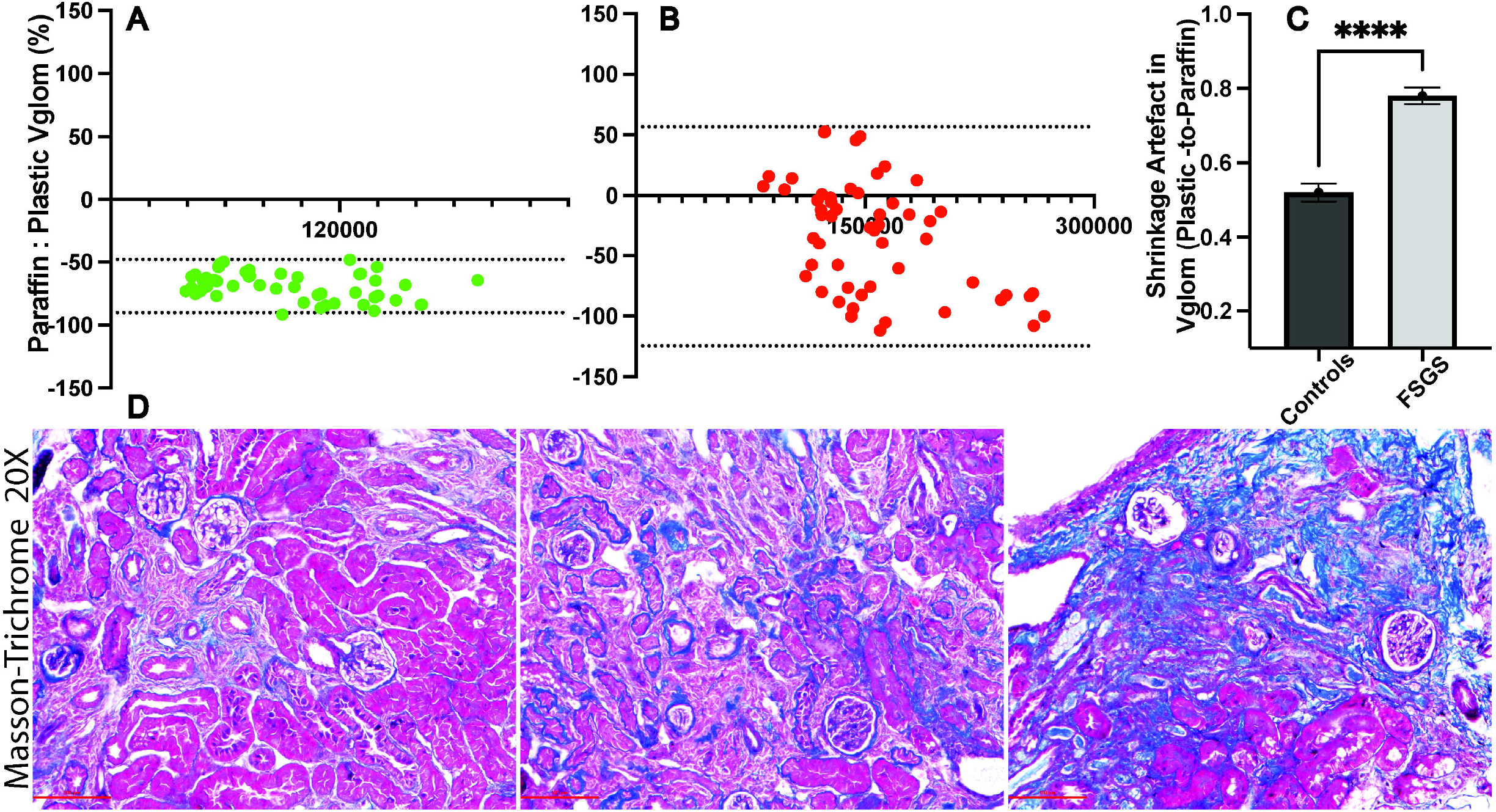
Differential shrinkage artefact and glomerular fibrosis in FSGS animals. Bland-Altman plots in **(A)** show ratios between paraffin:plastic measurements of MGV obtained in each control animal from 10- or 20-glomeruli *within* respective techniques [11 comparisons from 6 control animals (green dots)] while **(B)** shows *these same* ratios of MGVs obtained from 10-glomeruli in each FSGS animal [6 comparisons from 10 FSGS mice (red dots)]. **(C)** Bar graph compares the Shrinkage artefact between FSGS and control animals [Mean ±SEM; ****P<0.0001]. **(D)** Photomicrographs (20X) from kidney sections of representative FSGS animals within our dataset showing variable glomerular fibrosis (blue stain) in each.

We also recently reported increased MGV in mice models with FSGS compared to the respective controls in each model^10^. Here we re-evaluated this data given the contribution of reduced shrinkage artefact seen in FSGS glomeruli to the reported glomerulomegaly reported in FSGS. Table-3 compares MGVs in control and FSGS animals across plastic- and paraffin-based techniques used here. As expected, significant glomerulomegaly in FSGS was clearly identifiable in paraffin-based measurements (where control glomeruli would be expected to have greater reduction in Vglom and MGV). In plastic based methods, MGV was increased in FSGS animals vs controls, albeit insignificantly in many comparisons. Hence glomerulomegaly reported in paraffin-based morphometric studies of FSGS is likely a combination of true increases in Vglom, as well as a reflection of glomerular fibrosis and reduced shrinkage.

**Table-3:** Comparison of Mean glomerular volumes of FSGS animals vs Control animals using Plastic-based and Paraffin-based-measurements.

| Plastic |  | Control | FSGS | t-test |
| --- | --- | --- | --- | --- |
| | | Mean ± SD | Mean ± SD | p value |
| 20 Gloms | Cav-100X | 152372 ± 10065 | 190298 ± 59351 | 0.0631 |
| | Cav-40X | 150916 ± 10387 | 186639 ± 61062 | 0.0858 |
| 10 Gloms | Cav-100X | 160196 ± 17877 | 202079 ± 90207 | 0.167 |
| | Cav-40X | 159047 ± 18033 | 199747 ± 93145 | 0.1917 |
| | 2-profile-100X | 141827 ± 20381 | 190166 ± 82683 | 0.0895 |
| | 2-profile-40X | 142100 ± 20898 | 188322 ± 89160 | 0.1279 |
| | 3-profile-100X | 142087 ± 14929 | 205331 ± 74514 | **0.0169** |
| | 3-profile-40X | 141294 ± 17228 | 198864 ± 78478 | **0.036** |
| | WG-100X | 153452 ± 33993 | 191176 ± 86738 | 0.2166 |
| | WG-40X | 152511 ± 28765 | 192003 ± 88845 | 0.1978 |
| Paraffin |  | Control | FSGS | t-test |
| | | Mean ± SD | Mean ± SD | p value |
| 20 Gloms | 2-profile-40X | 70106 ± 10589 | 131679 ± 30657 | **0.0011** |
| | 3-profile-40X | 72824 ± 11679 | 133428 ± 32756 | **0.002** |
| | WG-40X | 78612 ± 14768 | 115469 ± 32886 | **0.037** |
| 10 Gloms | 2-profile-40X | 69527 ± 7062 | 120335 ± 35626 | **0.0092** |
| | 3-profile-40X | 71565 ± 4907 | 123480 ± 41251 | **0.0175** |
| | WG-40X | 81627 ± 20414 | 123880 ± 40305 | 0.0511 |

## METHODS

### Animals

We performed all our analyses in Balbc/J mice to avoid any strain-related variation in comparisons. We used young male adult BALBc/J as controls (2-6 months age) for evaluating baseline morphometry (n=10). To evaluate morphometry with significant glomerular injury, we used mice from two FSGS models which were also performed in BALBc/J background(n=10) - hypertrophic injury and a previously published ageing model^10,17^. The FSGS lesions in these mice models were confirmed and characterized by a renal pathologist [representative images in Supplementary Figure]. FSGS developed in hypertrophic 1/6^th^ remnant BALBc/J kidneys (after sequential 2/3^rd^ and contralateral nephrectomy) by 6 weeks ^15^. In the ageing model (mice > 12 months age), doxycycline feeding for 6 weeks, induced Shroom3 knockdown, diffuse podocyte foot process effacement and podocyte loss with the development of FSGS^10,17^.

### Tissue Preparation

<u>Plastic Embedded</u>: Kidneys were perfused with PBS cut into 1-mm cubes, post-fixed in glutaraldehyde, rinsed with buffer, post-fixed with 1% osmium tetroxide, dehydrated through an ethanol series and embedded in PolyBed 812. Sections were cut using a Reichert Ultracut E ultramicrotome set to cut 1-μm thick sections and stained with 1% toluidine blue. Section thickness was validated by embedding a section in PolyBed 812, cutting an ultrathin (silver-gold) section perpendicular to the original section plane, imaging the edge of the original section and measuring the thickness of the section. <u>Paraffin Embedded</u>: Kidneys were perfused with PBS at the end of the study in each animal. Representative lower pole tissue (where possible) was bisected and submerged in 4% formaldehyde followed by 70% ethanol. Paraffin-embedding and serial sectioning (every 5 microns x 5 sections) were done at Yale Pathology (Dr. Perincheri). Sections were labelled and stained with per-iodic acid Schiff (PAS) for Weibel-Gomez and 2/3-section Vglom measurements.

### Glomerular Volume Measurement Methods

#### Disector/Cavalieri (Plastic sections)

a. Sampling: A plastic block was arbitrarily selected, trimmed, faced and the first technically good section was saved to a slide, stained, coverglassed, and labeled slide 0. With the microtome continuous cutting every 10^th^ section was saved to a slide and sequentially numbered 10, 20, 30… until the 200^th^ section (21^st^ slide) was saved. Using the 10x lens of a Zeiss microscope, a map was drawn of section 0 and the location of all the glomerular profiles present in the section. Since an unknown volume of the glomeruli present in the stack of saved sections, glomeruli present in section is not usable. Next, section 10 is observed, and any profiles from newly appearing glomeruli are mapped, numbered sequentially from 1 and imaged using the 10X lens image. A digital camera attached to an Apple computer with 27” monitor used image glomeruli. After newly appearing glomeruli on section 10 have been mapped, numbered and imaged, section 20 is observed, new appearing glomeruli are mapped, numbered and profiles from all the numbered glomeruli are imaged. This is repeated until all profiles from the first 20 complete glomeruli have been imaged.
b. Measuring: Images were viewed using Adobe Photoshop, with window magnification of 67%. The Grid function of Photoshop was turned on with the grid lines 100 mm apart. Grid lines were red to contrast with the blue color of the sections. The magnification of the images was validated using a stage micrometer. The number of grid points falling on each profile from each of the 20 glomeruli was counted and logged. Volume of individual glomeruli is calculated with the equation

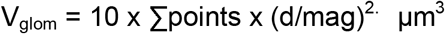

where 10 is the distance between sections in μm, Σpoints is the sum of grid points “hitting” all the profiles from a glomerulus, d is the distance between grid points in μm and mag is the magnification of the images.

#### 2-Profile method

The volume of a sphere can be determined from two arbitrary parallel profiles from the sphere a known distance apart. For this method glomeruli are assumed to be spheres and thus the two glomerular profiles are assumed to be circles. The areas of the two glomerular profiles were measured and the two radii calculated. The two radii can be used to calculate the radius of the sphere the two profiles came from ^13^.

For the 2-Profile method Sections 10, 20, 110, and 120 previously used for the Disector/Cavalieri method were imaged. Using the 10X lens of the microscope images of the complete section were taken from each of the four slides and used to identify profiles from individual glomeruli in the different sections. The first 20 glomeruli that intersected both Sections 10 and 20 or Sections 110 and 120 were selected and sequentially numbered for measurement [Figure 4]. The two profiles from each of the 20 glomeruli were imaged using both the 40X and 100X objective lenses. The same grid of points used for the Disector/Cavalieri method was used. The individual glomerular volumes (IGV_2-P_) were calculated using the equation:

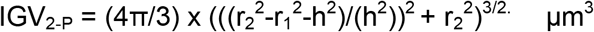

where r_1_ is the radius of the smaller glomerular profile, r_2_ is the radius of the larger glomerular profile and h is the distance in μm between sections.

Mean glomerular volume (MGV_2-P_) was determined by calculating the average of the glomeruli. Since only glomeruli that intersected the first section were used there may be a biased toward larger glomeruli. Also, the method assumes the glomeruli are spheres which they are not. This may also add a bias.

#### 3-profile method

We introduce the 3-Profile method here. To test whether using three profiles increases the precision of the individual glomerular volumes, sections 30 and 130 from the Disector/Cavalieri method were incorporated with the results of the 2-profile method. First the glomerular volume was calculated using the 2-profile method with the first and second sections then a second volume was calculated using the second and third sections. Finally, the individual glomerular volume (IGV3-P) was determined by calculating the average of the two 2-profile volumes. The MGV3-P for an animal was determined by finding the mean of the 10 or 20 glomeruli from the animal. The 3-Profile method will have similar biases as the 2-Profile method.

#### Weibel-Gomez method (for paraffin and plastic embedded tissue)

Sampling. Using the 100X lens image the profiles from all the glomeruli present in section 10. If necessary, use section 80 to image a total of 20 glomeruli.

Measurement: The Grid tool is turned on, with grid lines 100 μm apart, and the number of grid points “falling” on each glomerular profile are counted. Mean glomerular volume is calculated with the equation:

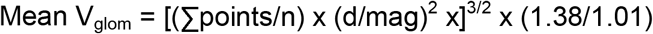

where Σpoints is the sum of grid points “hitting” all the glomerular profiles, n is the number of profiles measured, d is the distance between grid points, mag is the magnification of the images, 1.38 is the shape factor assuming the glomeruli are spheres, and 1.01 is correction factor assuming a coefficient of variation of 0.10 for the glomeruli within a kidney.

For paraffin-tissues in control animals we were able to get upto 50 glomerular sections in each animal. MGV using W-G in paraffin included 10-, 20- or 50-glomerular profiles. Larger glomeruli close to medulla or adjacent to large blood vessels were excluded where possible in all techniques.

#### Statistical Considerations

We assessed the concordance between gold-standard glomerular volume measurements and alternative measurements via Lin’s Concordance Correlation Coefficient^21^. This metric combines both precision and accuracy and increases (to a maximum of 1) as the line of best fit between two sets of measurements approaches the line of unity and as the observed data hews more tightly to that line. We used linear regression to determine the equation of a line that would best relate measurements from alternative assessments to the gold standard.

We also empirically tested the proportion of times sampling 10 random glomeruli would result in a mean volume within 5, 10, or 25% of the true mean, i.e., mean volume obtained by sampling 20 glomeruli. For this analysis, we randomly sampled 10 glomeruli from each mouse, calculated mean glomerular volume of the sample, and determined whether the mean volume of these sampled glomeruli was within 5, 10 or 25% of the true mean. We then repeated this random sampling procedure for a total of 1000 iterations and reported the proportion of times the mean obtained was within 5, 10 or 25% of the true mean. We report logit-transformed 95% confidence interval of this proportion by clustering at the level of the animal^22^. We present these data for all mice as well as based on whether the underlying diagnosis was FSGS or control. In this analysis we only included animals with 20 glomeruli sampled (=14). All analysis were conducted in STATA (version 14.2, College Station, TX).

## DISCUSSION

From experimental and clinical data, morphometric estimates of glomerular volume have diagnostic and prognostic implications, over and above information obtained from histology alone. The most precise technique for Vglom estimation Cavalieri/Fractionator-disector, is time-consuming to carry out, wastes tissue sections and requires considerable expertise limiting its utility in retrospective clinical studies. Here we used tissue samples, both plastic and paraffin, from control and FSGS mice to understand relationships between Vglom obtained by Cavalieri/Fractionator-disector technique and other reported techniques. In both paraffin and plastic, we also tested a novel 3-profile technique as a modification of the 2-profile method. Within the gold standard Vglom-Cav we also evaluated the precision of MGV estimation while using MGVs obtained from 10 IGV measurements vs 20 IGVs. In both FSGS and controls, we identified acceptable precision of sampling 10 but not 5 glomeruli such that only about 10% of Vglom-Cav with 10 glomeruli would be beyond 10% of 20-glomerular MGV, whereas almost 30% of Vglom-Cav with 5 glomeruli would show similar results. In plastic based techniques, 2- or 3-profile method measurements showed greater concordance with Vglom-Cav, than W-G method. The 3-profile technique offered incremental benefit to the existing 2-profile method. Changing image magnification (100X vs 40X) during points counting had minimum and inconsistent effect on MGV in the different techniques. Finally, we identified a consistent reduction of Vglom values within control animals (~50%) when using paraffin-based methods (vs corresponding methods in plastic embedded tissue) demonstrating a clear shrinkage artefact due to tissue processing. Most interestingly, FSGS glomeruli showed significantly less and more variable shrinkage artefact likely due to glomerular fibrosis.

Our findings have implications for experimental and clinical morphometric studies. Our data provides an estimate of precision when 10 glomeruli are randomly sampled and MGV is obtained using Vglom-Cav. This finding is consistent with prior reports ^13,18^. This number of glomerular profiles was present in the majority of clinical biopsies from a recent dataset from the NEPTUNE consortium^15^. Further, in spite of the interglomerular Vglom variability in FSGS, 10 glomerular MGV showed similar precision in both FSGS and controls. The less intensive 2- or 3-profile methods when performed in plastic also showed high concordance with Vglom-Cav in FSGS and controls, suggesting that despite biases (listed in Fig 1), these methods provided reasonable estimates of MGV when compared to Vglom-Cav, and could be used for animal or clinical studies. Our findings also clearly quantify the shrinkage artefact in MGV estimation when using paraffin-based methods. Hence estimations of MGV could utilize this “shrinkage factor” to calculate the true MGV. We believe this is especially applicable to animal data involving glomerular morphometry, especially if plastic-embedded samples were not obtained. This shrinkage factor would need to be reconsidered when significant and/or variable glomerular fibrosis is identified. In such cases, the paraffin based MGV estimates could indeed be suboptimal. On the other hand, the presence of less-than-expected shrinkage in paraffin could be a marker that signifies the presence of glomerular fibrosis.

We limited our analysis to BALBc mice (vs other backgrounds), which could be considered a limitation. Larger glomeruli close to the medulla or adjacent to large blood vessels were excluded where possible in all techniques although not universally. Our goal was to preferentially sample cortical glomeruli and avoid sampling larger Juxtaglomerular glomeruli. We acknowledge some variability in sample sizes due to the additional use of renal tissues, especially in FSGS animals, for other studies.

In summary, our detailed evaluation compares commonly applied glomerular morphometric techniques using both control and FSGS animals with plastic and paraffin tissue data. We quantify the precision of 5- or 10-glomerular MGV in either controls or FSGS and develop relating equations between different plastic-based measurements to the gold-standard Cavalieri method. We demonstrate clearly and quantify the shrinkage bias during tissue processing for paraffin-embedding. Finally, we report the role of diagnoses of FSGS and resultant glomerular fibrosis in altering the shrinkage artefact. Our findings have application to clinical and experimental studies using glomerular morphometry.

## Acknowledgements

MCM acknowledges research support from NIH RO1DK122164. FPW acknowledges research support from NIH R01DK11319 and R01HS027626. This research was further supported by P30DK079310. DGM is supported by K23DK117065 and R01DK128087

## Notes

### Competing Interest Statement

The authors have declared no competing interest.

